# Green-synthesized silver nanoparticles enhance *Guibourtia tessmannii* antithromboinflammatory therapeutic potential

**DOI:** 10.64898/2026.07.03.736249

**Authors:** Eya’ane Meva Francois, Gouli Lougui Linda Patricia, Nguemfo Edwige Laure, Fannang Simone Veronique, Ntoumba Agnes Antoinette, Bamal Hans-Denis, Thi Hai Yen Beglau, Tako Djimefo Alex Kevin, Mintang Fongang Ulrich Armel, Sone Enone Bertin, Armel Florian Tchangou Njiemou, Danielle Ines Madeleine Evouna, Yinyang Jacques, Chimi Tchatchouang Geordamie, Fonye Nyuyfoni Gildas, Janiak Christoph

## Abstract

**Introduction:** Thromboinflammation, which represents the pathological interplay between inflammation and thrombosis, is a leading cause of global mortality. Current therapies are frequently associated with an increased risk of bleeding and do not adequately address the inflammatory component of the disease. The African tree *Guibourtia tessmannii* represents a promising source of natural anti-inflammatory compounds. This study aimed to synthesize and characterize silver nanoparticles using an aqueous bark extract of *G. tessmannii* (GT-AgNPs) and to evaluate their anti-inflammatory and anticoagulant properties.

**Methods:** GT-AgNPs were synthesized by reducing silver nitrate with an aqueous extract of *G. tessmannii* bark. The nanoparticles were comprehensively characterized using UV-Vis spectroscopy, FTIR spectroscopy, powder X-ray diffraction, and scanning electron microscopy. In vitro anti-inflammatory activity was evaluated through inhibition of bovine serum albumin denaturation, whereas in vivo anti-inflammatory activity was assessed using the carrageenan-induced rat paw edema model. Anticoagulant activity was investigated by measuring activated partial thromboplastin time (aPTT) and prothrombin time (PT), corresponding to the intrinsic and extrinsic coagulation pathways, respectively.

**Results:** The synthesis successfully produced GT-AgNPs with an average particle size of approximately 20 nm. Both the aqueous extract and GT-AgNPs exhibited marked anti-inflammatory activity. The nanoparticles achieved 95% inhibition of protein denaturation in vitro and 95% inhibition of carrageenan-induced paw edema in vivo at a dose of 0.4 mg/kg body weight after 5 h. Furthermore, both the extract and GT-AgNPs demonstrated dose-dependent anticoagulant activity.

**Conclusion:** The study demonstrated that GT-AgNPs, synthesized from the bark of *G. tessmannii*, possess significant anti-inflammatory and anticoagulant properties. These findings highlight the potential of GT-AgNPs as nanotherapeutic candidates for the management of thrombo-inflammatory disorders.

## I. Introduction

Thrombosis is a leading and life-threatening complication of cardiovascular diseases, ranking among the top causes of death globally and posing a significant challenge for healthcare systems. Current antithrombotic strategies primarily target platelets and the coagulation cascade but are associated with a considerable risk of bleeding. Additionally, these treatments do not fully eliminate thrombotic events, suggesting a therapeutic gap likely linked to a third mechanism; the inflammation, that remains insufficiently addressed [1]. Analysis of data from 123 countries representing over 2.6 billion people revealed that pulmonary embolism (PE) accounted for 0.46% of total deaths (86,930 deaths) [2]. The analysis further disclosed that PE-related mortality increased with age and varied widely among countries (0–24 deaths per 100,000 person-years), with income level only partially explaining the observed disparities. Inflammation and thrombosis are closely linked through a host defense mechanism called immunothrombosis. When dysregulated, this process leads to thromboinflammation, characterized by excessive activation of platelets, innate immune cells, the complement system, and the coagulation cascade [3]. This contributes to micro- and macrovascular thrombosis and is associated with inflammatory conditions such as infections, autoimmune diseases, and clonal hematopoiesis [3]. Conventional antithrombotic therapies either prevent thrombus formation or dissolve existing thrombi but frequently produce severe adverse effects, including gastrointestinal hemorrhage and hemorrhagic stroke. Consequently, the search for safer therapeutic alternatives has increasingly focused on medicinal plants and their bioactive metabolites. Plants such as *Chamomilla recutita* L., *Allium sativum*, and *Rosmarinus officinalis* have demonstrated promising anticoagulant and antithrombotic activities through modulation of different components of the coagulation process [4, 5].

*Guibourtia tessmannii* is a semi-heliophilous, non-gregarious tree species native to Central Africa. It is commonly known as African rosewood or bubinga and is referred to locally as Essingang by the Ewondo people of Cameroon, Oveng or Fang in Equatorial Guinea, and Kevazingo in Gabon [6]. The tree can attain heights of up to 60 m and trunk diameters of approximately 2 m, with characteristic large sinuous buttresses [6]. Its bark is greenish-grey to reddish-brown, displaying numerous rounded scales that expose orange-red depressions beneath the surface [7]. In traditional medicine, *G. tessmannii* is used as an aphrodisiac, against witchcraft and evil spirits [8]. A decoction of its bark is prescribed for dysmenorrhea, cancer, high blood pressure, abortion prevention, malaria, scabies, itching, excessive white discharge, rheumatological pain, epilepsy, nervous disorders, various infections, gynecological disorders and diabetes [9–11]. Bark macerations are used in the treatment of headaches and dysentery, while bark powder is used in the treatment of colitis and against bewitchments [8, 9, 12].

Phytochemical studies have shown that the aqueous extract of *G. tessmannii* is rich in phenolic compounds including polyphenols, flavonoids, tannins and anthraquinones. Alkaloids and saponins have also been identified [13]. However, an absence of sterols, terpenes, reducing compounds and sometimes flavonoids has been noted [14]. Pharmacological studies have demonstrated a broad spectrum of biological activities, including cytotoxic, antioxidant [13,15,16], hypoglycemic, hypotensive [15], vasorelaxant [17], pro-ejaculatory [18,19], antimicrobial, and anti-inflammatory effects [20]. Furthermore, acute and subacute toxicity studies have shown that the aqueous bark extract is well tolerated, with no evidence of toxicity at oral doses up to 5000 mg/kg body weight [21,22].

Silver nanoparticles (AgNPs) possess unique optical, electronic, and antibacterial properties, which have led to their extensive application in fields such as biosensing, photonics, electronics, and antimicrobial therapies [23]. More recently, AgNPs have emerged as efficient nanocarriers for plant-derived bioactive compounds, improving their physicochemical stability, bioavailability, controlled release, and therapeutic efficacy. Such nanoformulations have shown considerable promise in antimicrobial, anti-inflammatory, antioxidant, and anticancer applications [24, 25]. The present study aimed to synthesize silver nanoparticles through the self-assembly of secondary metabolites present in the aqueous bark extract of G. tessmannii onto silver ions and to evaluate the anti-inflammatory and anticoagulant activities of the resulting phytogenic nanoparticles.

## II. Materials and Methodology

### II.1. Collection, authentication, and preparation of extract

The trunk bark of *Guibourtia tessmannii* was collected from Makounda village, located 12 km from Yabassi town and 7 km inside the forest toward Yinqui, in the Yabassi District, Nkam Division, Littoral Region of Cameroon. Botanical identification was carried out at the National Herbarium of Cameroon, referencing specimen No. 1481/SRF/Cam, and comparing it to the voucher material MPOM B. N°246. The bark was air-dried at room temperature (protected from direct sunlight) for four weeks, then grounded into a fine powder. For extraction, 20 g of the powdered material was mixed to 200 mL of distilled water preheated to 80 °C. The mixture was stirred continuously at 150 rpm for 5 min using a magnetic stirrer with a hot plate (BS4HC, Biobase, China). After cooling to room temperature, the extract was filtered using Whatman No. 1 filter paper. A portion of the filtrate was oven-dried at 40 °C for 24 hours to remove residual moisture. The dry extract was weighed, and the yield of extractable content was calculated according to the following formula (1):

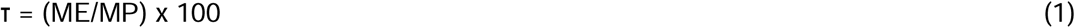

Where τ is the extraction yield; ME is the mass of the dried aqueous extract; and MP is the mass of the infused plant bark powder.

### II.2. Phytochemical screening

The aqueous extract of *G. tessmannii* stem bark was screened for the presence of major classes of secondary metabolites, including alkaloids, saponins, triterpenes, phenolic compounds, flavonoids, tannins, anthraquinones, and reducing sugars, using standard qualitative phytochemical procedures described previously [26].

### II.3. Green synthesis of silver nanoparticles from *G. tessmannii* aqueous extracts

A volume of 20 mL of aqueous plant extract (adjusted to pH 8) was added dropwise to 50 mL of 10^−2^DM silver nitrate (AgNO₃) solution under ambient conditions. The resulting mixture was incubated in the dark at room temperature for 7 days under static conditions to minimize photoactivation of AgNO₃ and allow for nanoparticle formation, as indicated by a visible color change. Following incubation, the colloidal suspension was centrifuged at 6000 rpm for 20 minutes using a Hettich centrifuge (D-7200 Tuttlingen, Germany). The resulting pellets were washed sequentially with deionized water and ethanol to remove unbound phytochemicals and residual ions. Purified nanoparticles were then dried in a hot air oven at 60D°C for 24 hours to obtain powders suitable for further characterization. The formation of silver nanoparticles was confirmed by UV-Visible spectrophotometry (Uviline 9100, Germany), with absorbance scanned over the range of 200–800 nm.

### II.4. Characterization of Biosynthesized AgNPs Fourier-transformed infrared spectroscopy (FTIR)

To investigate the role of phytochemicals in the reduction and stabilization of Ag^+^ into metallic Ag^0^ nanoparticles, FTIR spectra were recorded using a Bruker Tensor 37 spectrometer equipped with an attenuated total reflectance (ATR) accessory over the spectral range of 600-4000 cm⁻¹. The obtained spectra were used to identify the functional groups involved in the reduction and capping processes of the biosynthesized silver nanoparticles.

#### Powder X-ray diffraction (PXRD)

The PXRD diffraction of the nanoparticles was performed using a Bruker D2 Phaser powder diffractometer equipped with Cu Kα radiation (Kα₁ = 1.54060 Å; Kα₂ = 1.54443 Å; Kβ = 1.39225 Å). Samples were prepared as thin films on low-background silicon sample holders. The crystallite size of the synthesized GT-AgNPs was estimated from the diffraction data using the Scherrer equation (Equation 2):

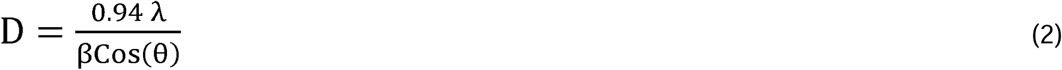

Where D is the average crystallite size (nm); λ is the X-ray wavelength (λ = 1.5406 Å); β is the Full width at Half Maximum (FWHM) of the diffraction peaks measured in radians; θ is the Bragg diffraction angle.

#### Scanning electron microscopy (SEM) and energy-dispersive X-ray spectroscopy (EDX)

The morphology of the nanoparticles was examined using a JEOL JSM-6510LV scanning electron microscope (SEM) equipped with a LaB₆ cathode operated at 20 kV and coupled to a Bruker XFlash 410 silicon drift detector for energy-dispersive X-ray (EDX) analysis (Bruker AXS, Karlsruhe, Germany). Prior to imaging, the nanoparticle samples were sputter-coated with gold using a JEOL JFC-1200 Fine Coater.

### II.5. Assessment of acute oral toxicity

The acute toxicity of the synthesized nanoparticles was assessed according to OECD guidelines 425 [24] at a limit dose of 2000 mg/kg. Female rats, 8-12 weeks of age and weighing 120-200 kg, were divided in 2 batches (n = 3). The rats were fasted 24 h before the test. One batch was administered the GT-AgNPs orally at the abovementioned limit dose, while the other received 10 mL/kg of distilled water for control. The animals were subsequently observed individually during the first 30 min following administration, then regularly during the first 24 h, and afterwards, once daily for 14 days. Parameters of interest were the aspect of the skin, hair, mucous membranes, eyes and the behaviour (tremors, convulsions, mobility, aggressive behaviour and lethargy). Body weight was recorded every 2 days. On day 14, the rats were humanly sacrificed under anaesthesia and the blood was collected after incision of the carotid artery for the analysis of renal (urea/creatinine) and hepatic (AST/ALT) biochemical parameters. The mass of vital organs (liver, kidneys, lung, heart and spleen) was also recorded.

### II.6. Experimental animals

Healthy adult female Wistar albino rats weighing between 120 - 200 g were obtained from the Laboratory Animal Centre of the Department of Pharmaceutical Science, Faculty of Medicine and Pharmaceutical Science, University of Douala, Cameroon. The animals were randomly sorted in standard polypropylene cages in groups of three and maintained under standard conditions of temperature (24 ± 2 °C) and light (approximately 12 h/12 h light/dark cycle) with free access to standard laboratory diet and tap water *ad libitum*.

### II.7. Heat-induced egg albumin denaturation assay

The anti-inflammatory activity of silver nanoparticles with *G. tessmannii* extract was assessed *in vitro* using the heat-induced albumin denaturation assay, as previously described by Chandra *et al.* [28]. The reaction mixture consisted of 0.2 mL of fresh egg albumin collected from chicken eggs and less than 24 hours old, 2.8 mL of phosphate-buffered saline (PBS, pH 6.4), and 2 mL of our nanoparticles and aqueous extract solutions prepared at concentrations series of 25, 50, 100, 200, and 400 µg/mL. For the standard, Diclofenac (2 ml) was used and prepared in the same concentration series, while distilled water (2 ml) was used for control. The reaction mixtures were incubated at 37 °C for 15 minutes, then gradually heated up to 70°C and kept at that temperature for 5 min before allowing them to cool down for 20 min. The tests were ran in triplicate. The absorbance of each reaction mixture was then measured with a spectrophotometer and the reaction percentage was calculated using the following formula (3):

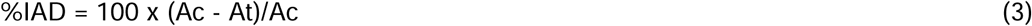

where IAD is the percentage inhibition of albumin denaturation, Ac is the absorbance of the control, and At is the absorbance of the test sample.

### II.8. *In vivo* anti-inflammatory assay

The *in vivo* anti-inflammatory effect of the silver nanoparticles was evaluated using the carrageenan-induced paw edema model in Wistar rats, as described by Winter *et al.* [29]. Five groups (n = 5) of rats were formed and fasted for 12 hours prior to the experiment. For each rat, the initial volume (V₀) of the left hind paw was measured using a plethysmometer. The different solutions were administered orally. Three groups received the silver nanoparticles with extract at 100, 200 and 400 μg/kg, respectively. One group received Diclofenac (10 mg/kg) and served as positive control. The last group received distilled water (10 ml/kg) and served as negative control. An acute inflammation was induced 30 min after administration of the solutions, by a single subplantar injection of 0.1% carrageenan (1% carrageenan suspended in 0.9% NaCl) into the left hind paw of the animals. A plethysmometer was used to measure the anti-inflammatory effect by measuring the volume of the left hind paw of each rat before and after administering the different solutions, at 30 min, then every hour post injection, for 6 h. The percentage of inhibition was obtained using the following formula (4):

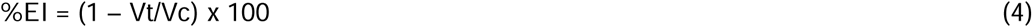

where EI is the percentage inhibition of edema, Vt is the mean edema volume of the treated group, and Vc is the mean edema volume of the control group.

### II.9. Preparation of erythrocyte suspension

An erythrocyte suspension was prepared following a protocol described by other authors, with slight modifications [27]. Rats’ whole blood (5 mL) was collected into EDTA tubes and centrifuged at 3000 rpm for 10 minutes. After discarding the supernatant, the red cell pellet underwent three successive washes with isotonic physiological saline (0.9% NaCl) until the supernatant became clear. The volume of packed cells was measured, and the pellet was resuspended in physiological saline to a final concentration of 10% (v/v), forming the red blood cell suspension used for *in vitro* hemolysis assays.

### II.10. Evaluation of hemolytic activity

The hemolytic activity of the tested aqueous extracts and silver nanoparticles was evaluated using a modified protocol based on Ouattar *et al.*, 2022 [27]. In 2 mL centrifuge tubes, 20 μL of each sample (nanoparticles or plant extract) at concentrations of 25, 50, 100, 200, and 400 μg/mL was mixed with 1980 μL of the erythrocyte suspension. The mixtures were incubated in a water bath at 37 °C for 60 minutes. After incubation, 250 μL of each reaction mixture was transferred into 750 μL PBS, immediately cooled in an ice bath to stop the reaction, and centrifuged at 3000 rpm for 10 minutes to sediment the intact erythrocytes. Under the same conditions and the same experimental procedures, a total hemolysis tube was prepared with 100 μL of the erythrocyte suspension and 1900 μL of distilled water for positive control, and another tube with 250 μL of the erythrocyte suspension and 750 μL of PBS for negative control. The absorbance of the supernatants was measured at 548 nm using a UV-Visible spectrophotometer (Uviline 9100, Germany), with PBS as the blank reference.The hemolysis rate of different solutions was calculated in relation to the total hemolysis, after 60 min of incubation according to the following formula (5):

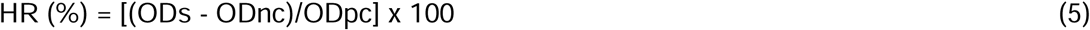

where HR is the hemolysis rate, ODs is the optical density of the sample, ODnc is the optical density of the negative control, and ODpc is the optical density of the positive control.

### II.11. Preparation of platelet-poor plasma

Platelet-poor plasma (PPP) was prepared according to the protocol of Atukorala *et al*. [30]. Whole blood was collected from the carotid artery of four Wistar rats into tubes containing 3.2% sodium citrate (blood:sodium citrate ration 9:1) and centrifuged at 3000 rpm for 10 min.The resulting PPP was collected and stored at –4D°C until further use.

### II.12. Activated partial thromboplastin time (aPTT)

Activated partial thromboplastin time (aPTT) was determined to evaluate the anticoagulant activity of the samples through the intrinsic coagulation pathway, following previously reported methods [30, 31]. Briefly, a Kaolin reagent kit was used to constitute a partial thromboplastin reagent (Kaolin Platelet substitute mixture), with cephalin as phospholipid substitute, and was prepared as per manufacturer’s guidelines, then pre-warmed at 37°C, in a water bath, together with a 0.025 M calcium chloride solution. The activity of the nanoparticles and extracts was carried out in clotting tubes containing 90 µL of PPP mixed with 10 µL of our GT-AgNPs samples at different concentrations (25, 50, 100, 200, 400 µg/mL) and 10 μL of PBS for control. After 15 min of incubation at 37°C (3 min for control), 100 μL of cephalin-kaolin was added to the mixture and further incubated for exactly 3 min at 37°C. The coagulation time was then determined using a stopwatch started after adding 100 μL of pre-warmed calcium chloride at 0.025 M. Enoxaparin was used as standard at 0.2 µg/ml.

### II.13. Prothrombin time (PT)

The prothrombin time allowed to assess the anticoagulant activity with respect to the exogenous coagulation pathway (factors VII, X, V, II, and fibrinogen), following previously reported methods [30, 31]. PPP (90 μl) preheated for 2 min at 37°C was mixed with 10 μL of our test solutions (*G. tessmannii* aqueous extract and silver nanoparticles with *G. tessmannii* extract) at different concentrations (25, 50, 100, 200, 400 μg/mL). After 15 min of incubation at 37°C, 200 μL of calcium thromboplastin (preheated for at least 15 minutes at 37°C) was added to the mixture and the coagulation time is then recorded using a stopwatch. Enoxaparin was used as standard at 0.2 µg/mL.

### II.14. Statistical analysis

Quantitative data from all tests were recorded and treated on MS Excel 2016 and analyzed using GraphPad Prism 10.5.1 for Windows (San Diego, California, USA). Results were presented as meanD±Dstandard error of the mean (SEM). Statistical comparisons were performed using two-way analysis of variance (ANOVA) followed by Tukey’s multiple comparison test. Differences were considered statistically significant at p < 0.05. Characterization graphs and crystallite size calculations were generated using OriginPro version 6.0 (OriginLab Corporation, Northampton, MA, USA).

## III. Results

### III.1. Extraction yield of *Guibourtia tessmannii* stem bark

Twenty grams of powdered *Guibourtia tessmannii* stem bark were extracted with 200 mL of distilled water. The resulting infusion was filtered and dried, yielding 1.5 g of a dark-brown, viscous residue. The extraction yield was calculated to be 7.5%.

### III.2. Phytochemical composition of the extract

Qualitative phytochemical screening of the aqueous extract of *G. tessmannii* stem bark revealed the presence of several classes of secondary metabolites, including phenolic compounds (flavonoids, tannins, and anthraquinones) and terpenoids (saponins). In contrast, coumarins, sterols, alkaloids, and reducing sugars were not detected (Table 1).

**Table 1.** Phytochemical content of the aqueous extract of *Guibourtia tessmannii* stem bark.

| Tested metabolites | Results |
| --- | --- |
| Alkaloids | - |
| Flavonoids | + |
| Tannins | + |
| Anthraquinones | + |
| Coumarins | - |
| Saponins | + |
| Sterols and terpenes | - |
| Reducing sugars | - |

(+) presence ; (-) absence

### III.3. UV-Visible spectroscopy of synthesized silver nanoparticles

The formation of GT-AgNPs was initially indicated by a visible color change of the reaction mixture from light brown to dark brown, confirming nanoparticle formation within the reaction medium (Figure 1).

**Figure 1.**
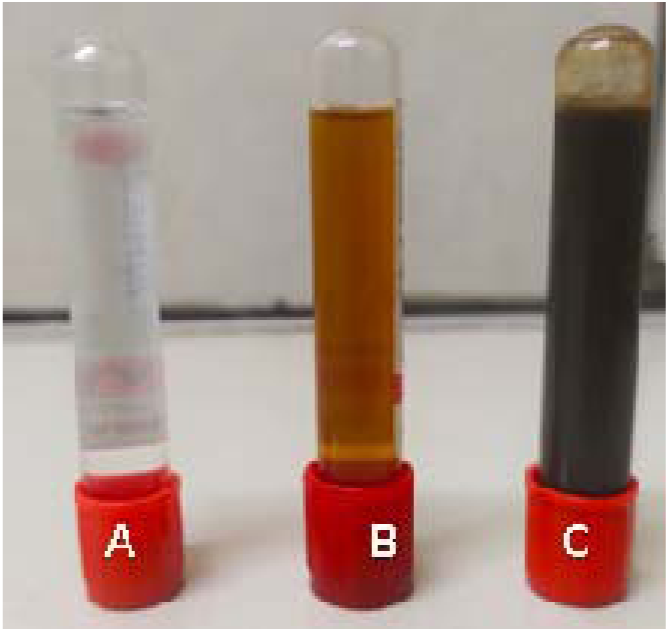
Color change observed during the synthesis of GT-AgNPs. (A) Silver nitrate solution; (B) aqueous extract of G. tessmannii; (C) mixture of silver nitrate and G. tessmannii aqueous extract.

The synthesis of the nanoparticles was further investigated using UV-Vis spectroscopy, which revealed a surface plasmon resonance band between 350 and 500 nm, with a characteristic absorption peak centered at approximately 450 nm and a maximum absorbance of 1.8, confirming the formation of silver nanoparticles (Figure 2). The UV-Vis spectrum of the nanoparticles obtained was compared with those of the aqueous extract and the silver nitrate solution.

**Figure 2.**
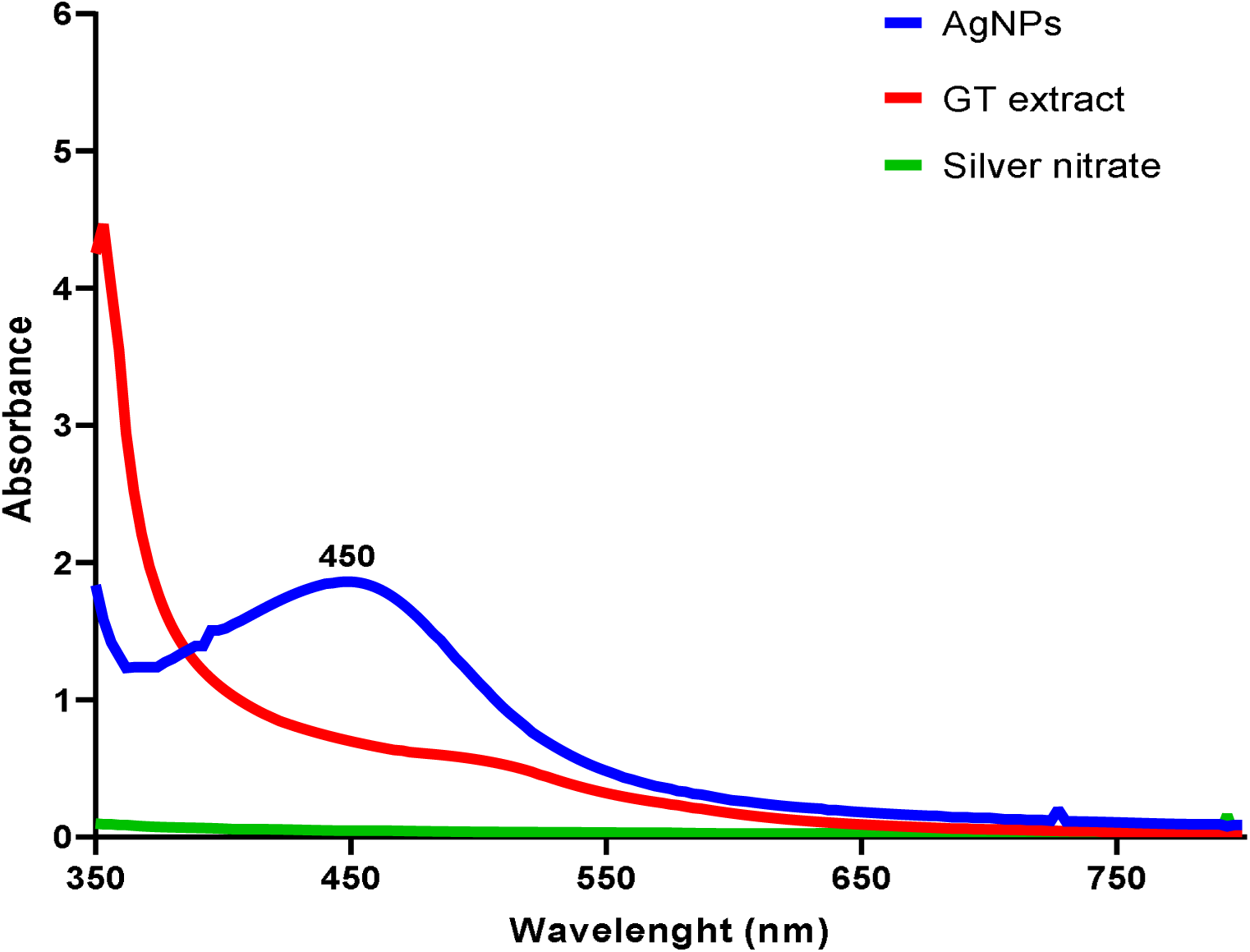
UV-Visible spectra of GT-AgNPs, aqueous *G. tessmannii* extract and AgNO_3_.

### III.4. Powder X-ray diffraction (PXRD)

PXRD analysis on green-synthesized GT-AgNPs revealed their crystalline nature. The diffractogram confirmed the nanocrystalline structure of the synthesized nanoparticles (Figure 3).

**Figure 3.**
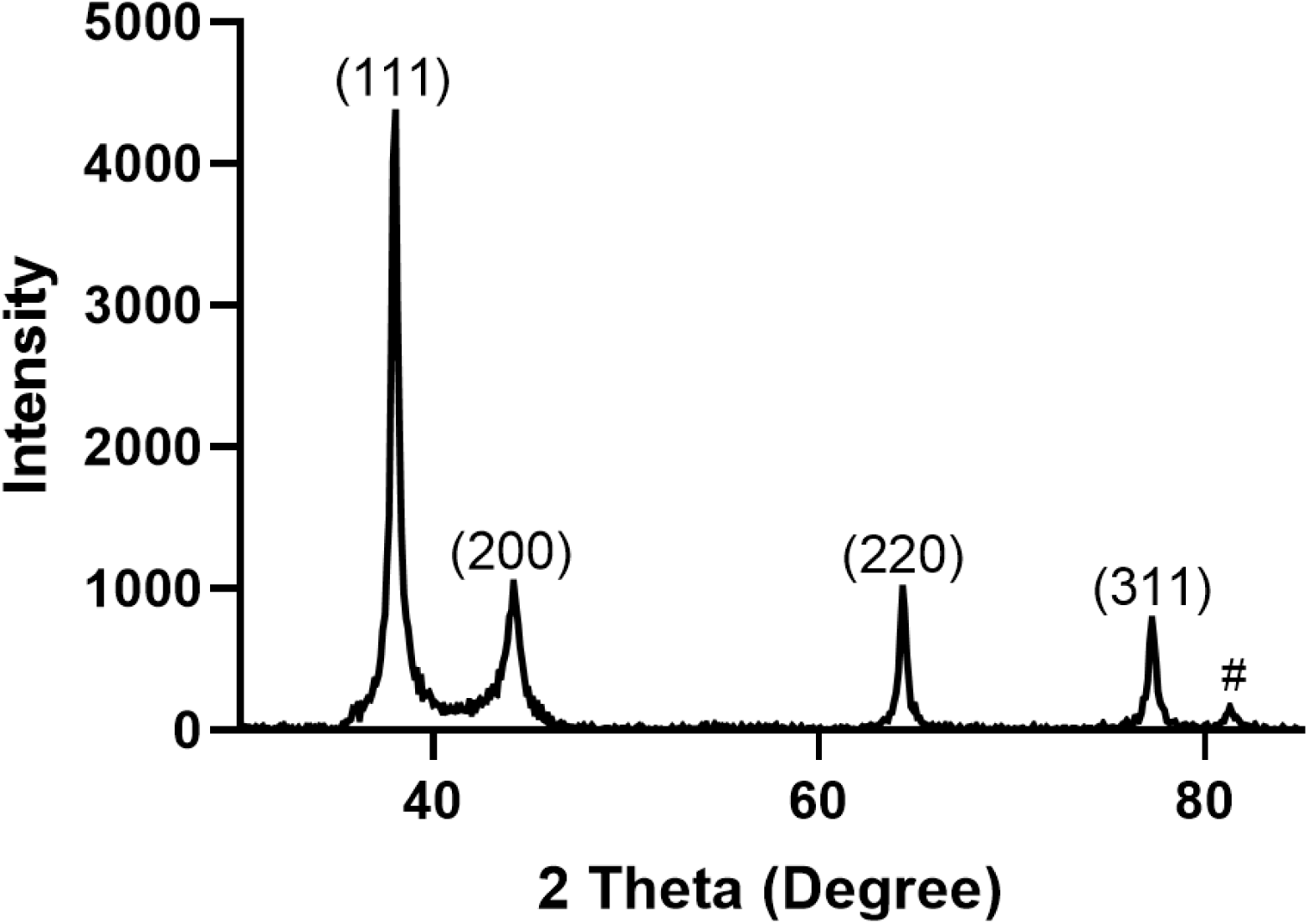
PXRD diffractogram of synthesized GT-AgNPs.

The diffractogram, recorded over a 2θ range of 30° to 85°, displayed four characteristic diffraction peaks at 38.01°, 44.16°, 64.34°, and 77.25°, corresponding to the (111), (200), (220), and (311) crystallographic planes of the face-centered cubic (fcc) structure of metallic silver, with a lattice parameter of 4.08 Å. These reflections were in agreement with the ICDD reference pattern No. 04-0783, confirming the crystalline nature of the synthesized nanoparticles. Application of the Scherrer equation to the diffraction data yielded an average crystallite size of approximately 20 nm, further confirming the successful biosynthesis of silver nanoparticles.

**Table 2.**
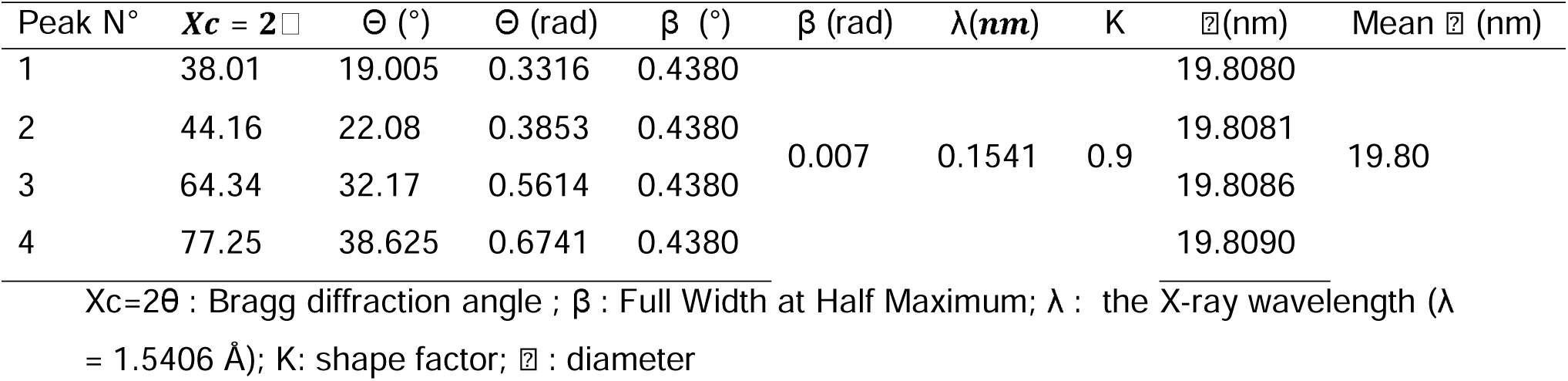
Crystallographic parameters of GT-AgNPs.

### III.5. Fourier Transform Infrared Spectroscopy (FTIR)

The FTIR of GT-AgNPs and *G. tessmannii* are depicted in figure 4. Several similarities were observed between the FTIR spectra of GT-AgNPs and the aqueous extract of *G. tessmannii* (Figure 4). The principal absorption bands of the plant extract appeared at 3358, 2928, 1632, 1436, 1346, 1248, 1022, 562, 517, and 411 cm⁻¹. In contrast, the FTIR spectrum of GT-AgNPs exhibited prominent bands at 3328, 2936, 1753, 1632, 1504, 1444, 1421, 1376, 1323, 1240, and 1044 cm⁻¹.

**Figure 4.**
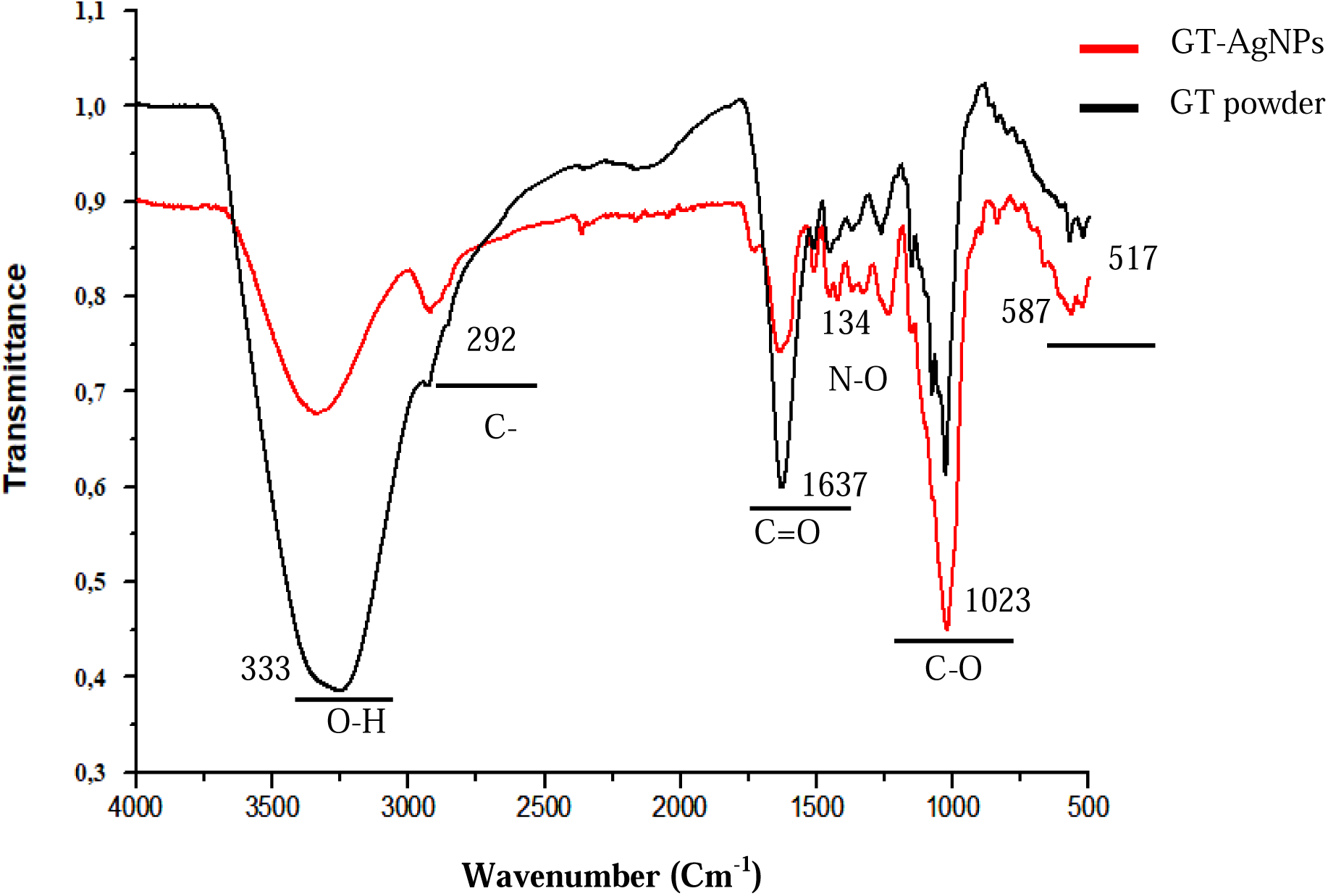
FTIR spectra of GT-AgNPs and *G. tessmannii* powder.

According to standard infrared band assignments, the broad absorption band around 3328 cm⁻¹ corresponds to the O–H stretching vibration. Compared with the plant extract, this band exhibited a red shift of approximately 20 cm⁻¹, suggesting the involvement of hydroxyl groups in the reduction and stabilization of silver nanoparticles. New absorption bands observed at 1736, 1421, and 1376 cm⁻¹ were assigned to carbonyl (C=O) and aromatic C=C stretching vibrations, indicating possible interactions between phytochemicals and the nanoparticle surface. The remaining absorption bands corresponded to stretching, bending, and deformation vibrations characteristic of functional groups naturally present in *G. tessmannii*, suggesting that these biomolecules acted as both reducing and capping agents during nanoparticle biosynthesis.

### III.6. Scanning electron microscopy (SEM) and energy dispersive X-ray Spectroscopy (EDX)

The SEM micrograph (Figure 5) of *Guibourtia tessmannii*-mediated silver nanoparticles (GT-AgNPs) provided visual evidence of their morphology and approximate size. The biosynthesized nanoparticles appeared predominantly as aggregated structures with irregular fibrous morphologies.

**Figure 5.**
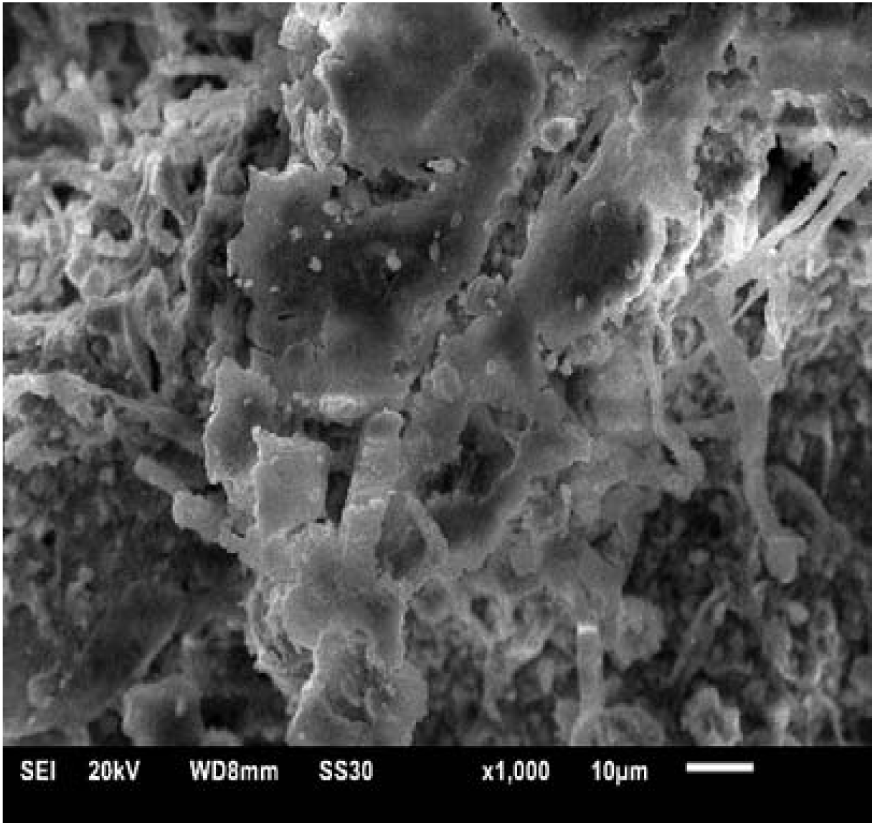
SEM image of of *Guibouria tessmannii*-mediated silver nanoparticles

Furthermore, energy-dispersive X-ray spectroscopy (EDX) (Figure 6) confirmed the elemental composition of the synthesized nanoparticles, revealing the presence of carbon, oxygen, and metallic silver. A strong characteristic silver signal was observed at approximately 3.0 keV, confirming the successful formation of silver nanoparticles.

**Figure 6.**
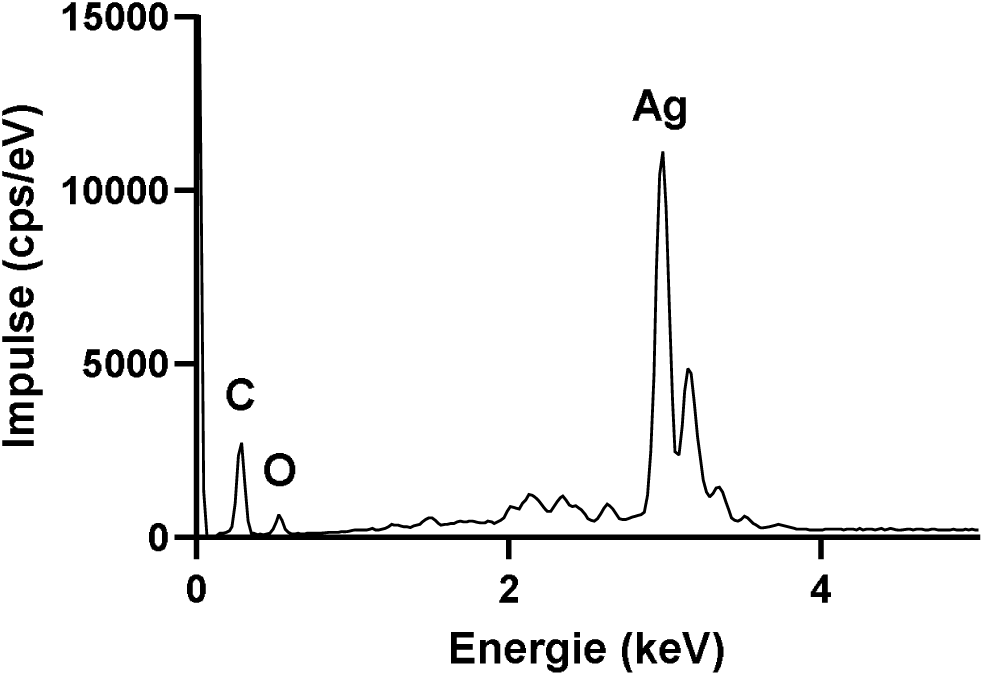
Energy dispersive X-ray spectrum of GT-AgNPs

### III.7. Acute oral toxicity

The acute toxicity of silver nanoparticles synthesized from G. tessmannii bark extract (GT-AgNPs) was evaluated according to OECD Guideline 425 using a limit dose of 2000 mg/kg body weight over a 14-day observation period. No mortality or treatment-related clinical abnormalities were observed, and all monitored parameters remained within normal physiological ranges (Table 3).

**Table 3.**
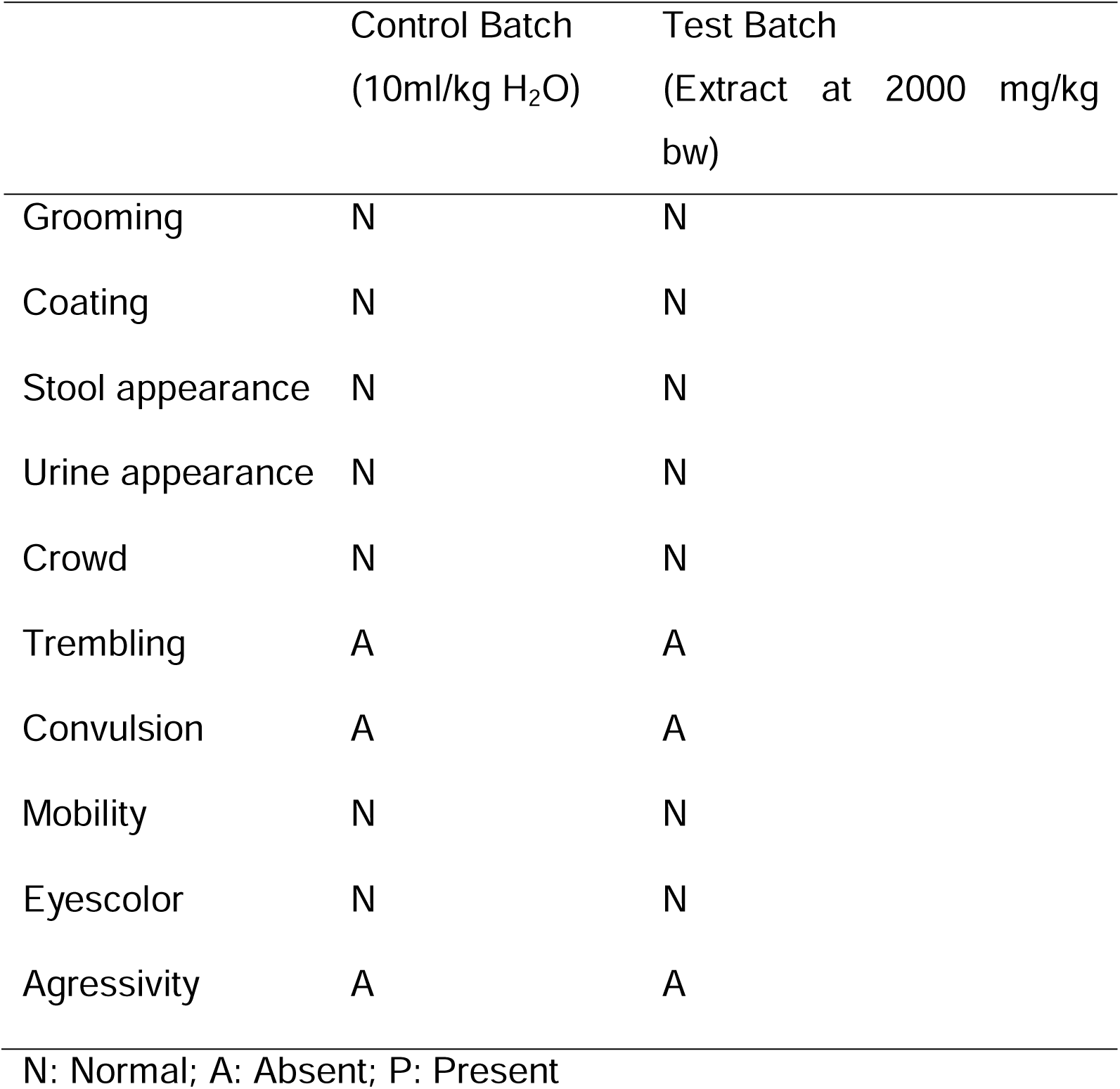
Clinical parameters observed during 14 days.

Furthermore, the rats were weighed for 14 days and their growth rates were calculated. The analysis revealed a normal increase in the weight of the test batch over a 14 days period (up to 29±3 g) and no significant difference was observed against the control batch (up to 25±2 g) despite the test batch showing slightly higher growth rate (Figure 7).

**Figure 7.**
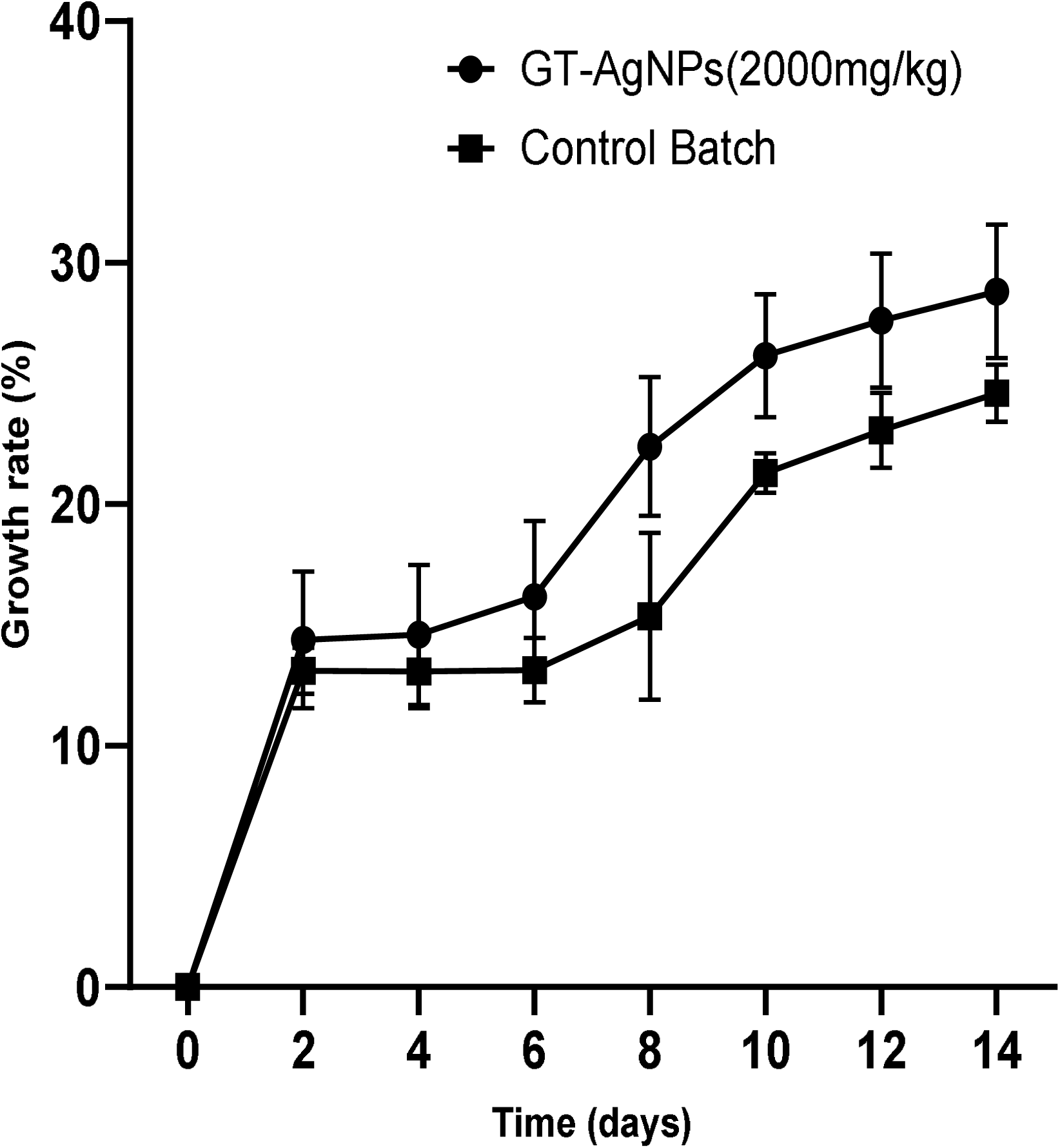
Time-course of body weight changes in rats treated with GT-AgNPs. Results are presented as mean ± SEM (n = 3). No statistically significant differences were observed between groups (p > 0.05).

Similarly, no significant differences were observed between the treated and control groups regarding serum creatinine and urea concentrations (Figures 8A and 8B, respectively), alanine aminotransferase (ALT) activity, or the relative weights of vital organs (Figure 8D). However, serum aspartate aminotransferase (AST) activity was significantly increased (p < 0.05) in the GT-AgNP-treated group compared with controls (Figure 8C).

**Figure 8.**
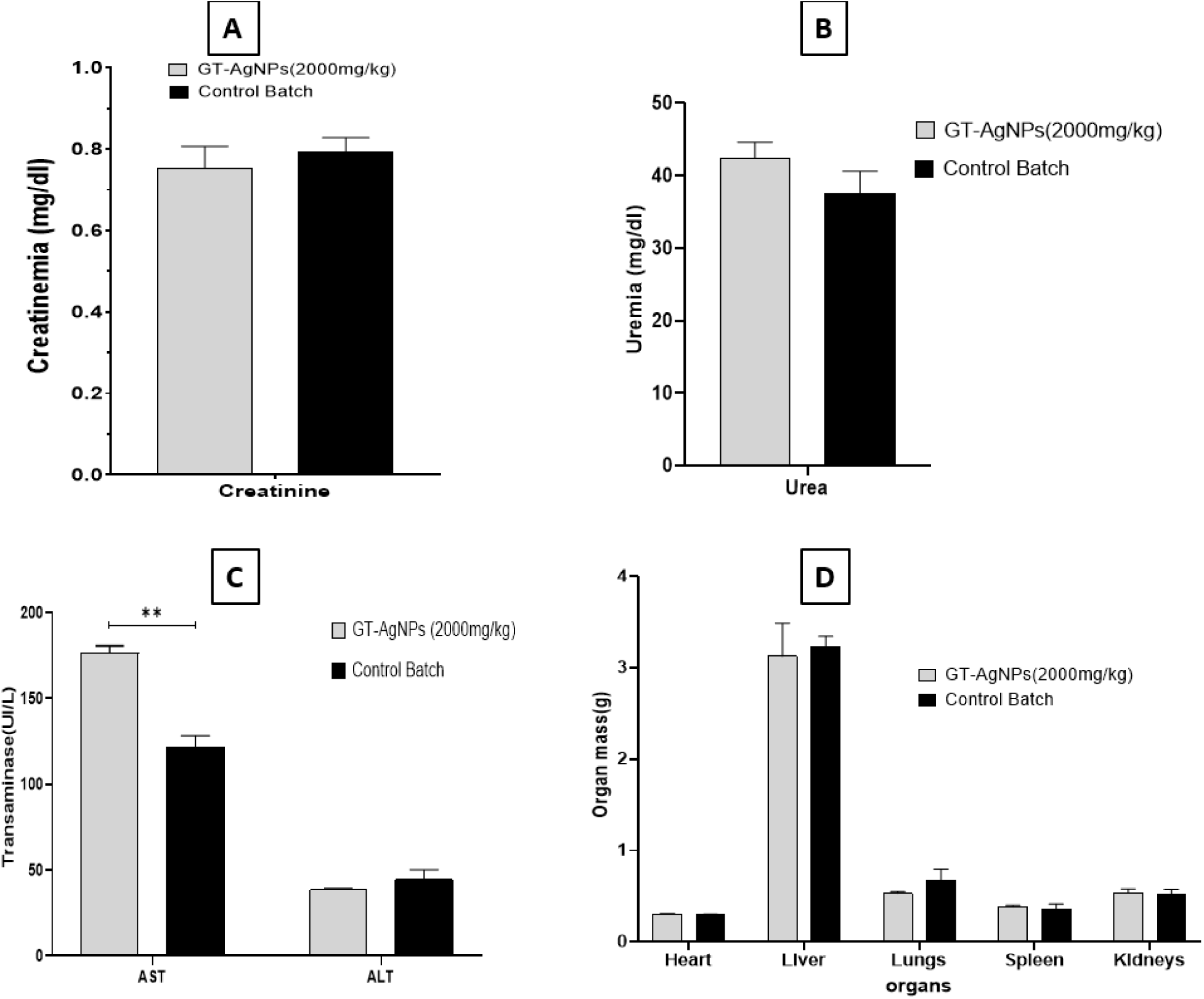
Effect of acute exposure to *Guibourtia tessmannii*-mediated silver nanoparticles on physiological parameters. (A) Serum creatinine; (B) serum urea; (C) hepatic enzymes (ALT and AST); (D) relative organ-to-body weight ratios. Results are presented as mean ± SEM (n = 3). *p < 0.05 versus control.

### III.8. Effect on erythrocytes hemolysis

The hemolysis rates observed after incubation of erythrocytes with the aqueous extract and GT-AgNPs are presented in Figure 9. Both the aqueous extract and GT-AgNPs exhibited low, dose-dependent hemolytic activity, with hemolysis ranging from 5–10% and 3–9%, respectively. Distilled water, used as the positive control, induced 100% hemolysis. According to accepted hemocompatibility criteria, hemolysis values below 10% indicate that both preparations are non-hemolytic.

**Figure 9.**
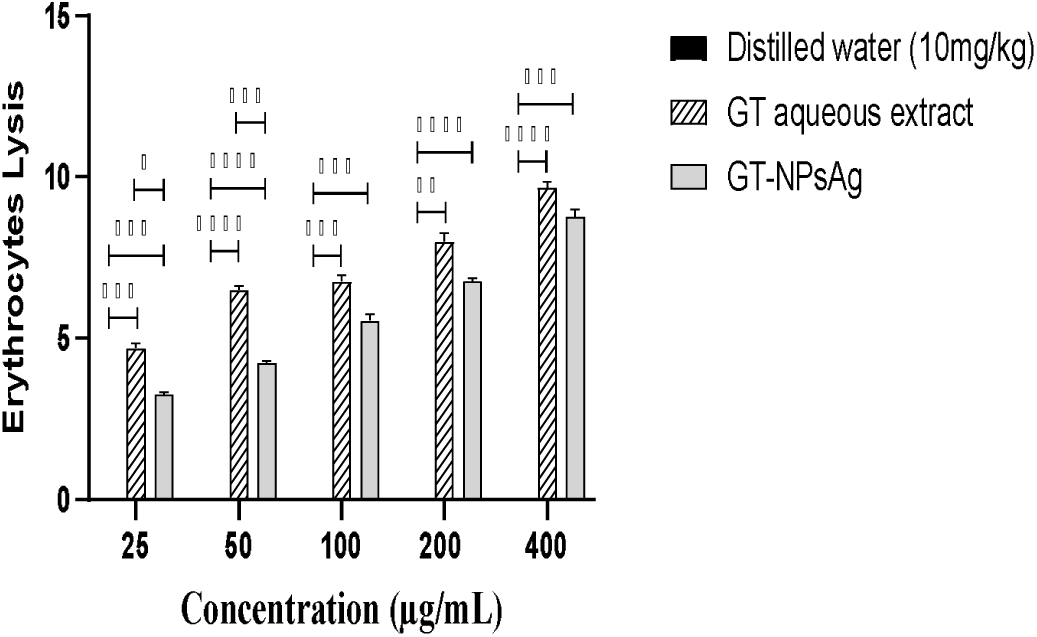
Evaluation of erythrocyte hemolysis.

### III.9. *In vitro* anti-inflammatory activity

The in vitro anti-inflammatory activity of the different samples, assessed by heat-induced egg albumin denaturation, is presented in Figure 10. All samples exhibited significant, dose-dependent inhibition of albumin denaturation. The highest inhibition was observed at 400 μg/mL, reaching 95% for GT-AgNPs and 87% for the aqueous extract. Under the same conditions, diclofenac, used as the reference anti-inflammatory drug, produced 70% inhibition.

**Figure 10.**
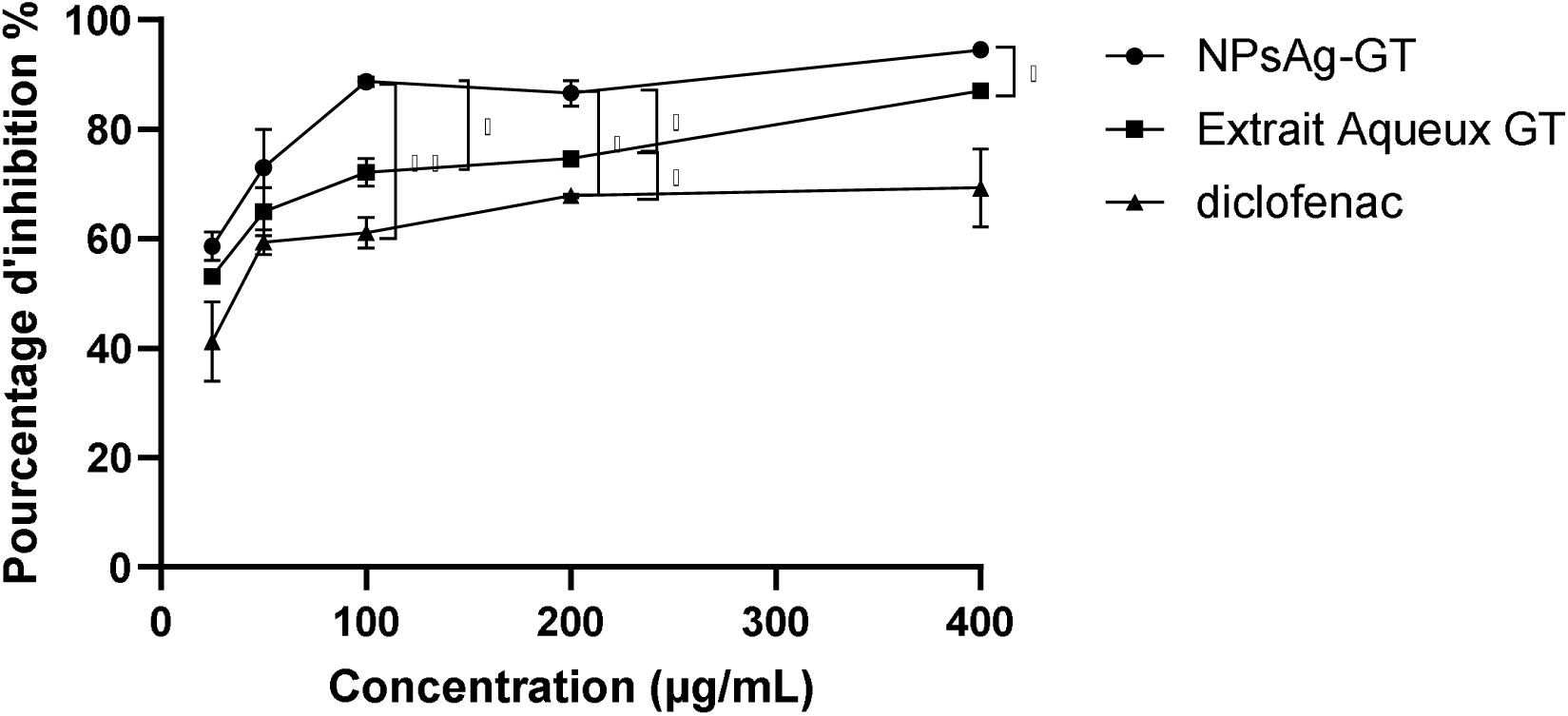
Inhibition of heat-induced egg albumin denaturation.

### III.10. Effect of the treatments on carrageenan-induced rat paw edema

The effects of the different treatments on carrageenan-induced paw edema are summarized in Table 4, together with their corresponding percentages of edema inhibition. Carrageenan injection induced progressive paw edema, which became evident within 30 minutes and reached its maximum intensity at 1 hour in the treated groups.

**Table 4.** Effect of the different treatments on carrageenan-induced paw edema. The maximum edema volume (highlighted in gray) occurred 1 hour after carrageenan injection in all treated groups. All treatments produced significant inhibition of edema compared with the control group.

| Groups | Dose<br>mg/kg | Measured paw volumes in mL (percentage of inhibition) |  |  |  |  |  |  |
| --- | --- | --- | --- | --- | --- | --- | --- | --- |
|  |  | 0.5h | 1h | 2h | 3h | 4h | 5h | 6h |
| **Distilled Water** | 10 | 1.4±0.1 | 2.2±0.2 | 2.2±0.2 | 2.7±0.1 | 2.7±0.1 | 2.9±0.1 | 2.8±0.1 |
| **Diclofenac** | 10 | 1.2±0.3<br>(15%) | 1.6±0.2a<br>(38%) | 1.6±0.2a<br>(39%) | 1.5±0.2c<br>(64%) | 1.5±0.2d<br>(68%) | 1.6±0.2d<br>(62%) | 1.7±0.1d<br>(55%) |
| **GT-AgNPs** | 0.1 | 1.12±0.09a<br>(14%) | 1.4±0.1b<br>(45%) | 1.3±0.1c<br>(62%) | 1.30±0.09d<br>(75%) | 1.3±0.1d<br>(78%) | 1.2±0.1d<br>(82%) | 1.3±0.1d<br>(79%) |
| **GT-AgNPs** | 0.2 | 1.13±0.08b<br>(31%) | 1.68±0.09b<br>(25%) | 1.54±0.2b<br>(44,89%) | 1.4±0.1d<br>(74%) | 1.2±0.1d<br>(86%) | 1.11±0.08d<br>(91%) | 1.3±0.1d<br>(80%) |
| **GT-AgNPs** | 0.4 | 1.2±0.1a<br>(18%) | 1.8±0.1a<br>(12%) | 1.6±0.1b<br>(42%) | 1.40±0.03d<br>(71%) | 1.21±0.12d<br>(85%) | 1.05±0.06d<br>(95%) | 1.24±0.07d<br>(83%) |

a= p<0.05; b =p<0.01; c= p<0.001; d=p<0.0001.

Oral administration of GT-AgNPs produced significant inhibition of paw edema compared with the control group. The maximum inhibition percentages observed at 5 hours were 82%, 91%, and 95% following doses of 0.1, 0.2, and 0.4 mg/kg body weight, respectively. By comparison, diclofenac sodium, used as the reference drug, achieved 68% inhibition at 4 hours.

### III.11. Evaluation of the intrinsic coagulation pathway (aPTT)

The anticoagulant activity of the aqueous extract and GT-AgNPs on the endogenous coagulation pathway was evaluated by measuring the activated partial thromboplastin time (aPTT) (Figure 11). Prolongation of aPTT relative to the control (normal range: approximately 30–34 s, depending on the reagent used) indicates inhibition of the intrinsic coagulation pathway.

**Figure 11.**
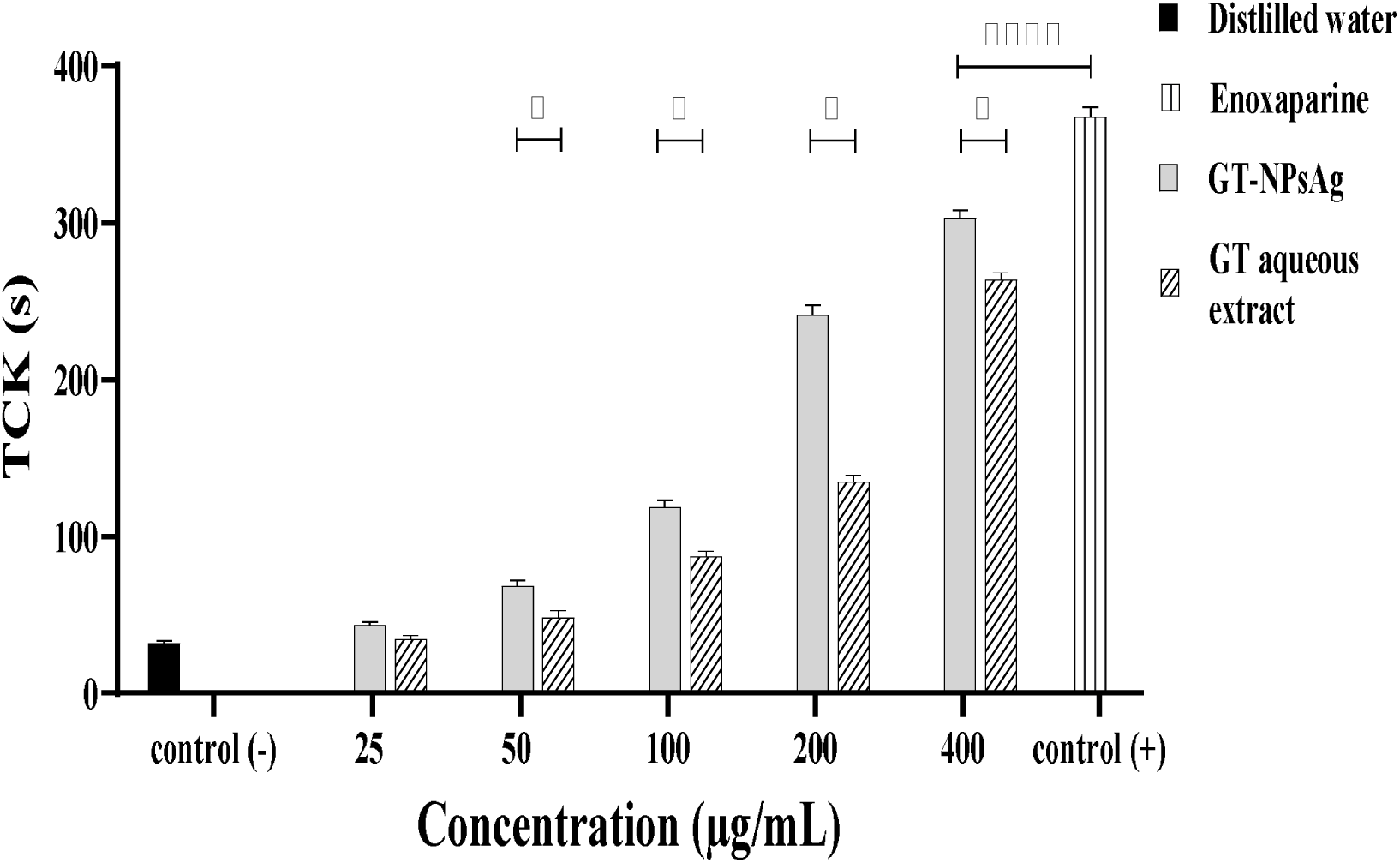
Evaluation of anticoagulant activity with respect to the endogenous pathway.

Both the aqueous extract and GT-AgNPs produced dose-dependent prolongation of aPTT, demonstrating anticoagulant activity. However, their effects remained lower than that of enoxaparin (0.2 mg/mL), which prolonged coagulation time to more than 300 seconds.

At the highest concentration tested (400 μg/mL), GT-AgNPs prolonged aPTT to 298 s, representing approximately a nine-fold increase compared with the negative control (32 s) and an extension of 266 s. Under the same conditions, the aqueous extract prolonged aPTT to 257 s, corresponding to approximately an eight-fold increase over the control and a prolongation of 225 s.

### III.12. Exploration of the exogenous pathway

The prothrombin time (PT) of normal plasma typically ranges between 12 and 14 seconds. Therefore, any prolongation of PT relative to the negative control indicates inhibition of the exogenous coagulation pathway. The corresponding results are presented in Figure 12.

**Figure 12.**
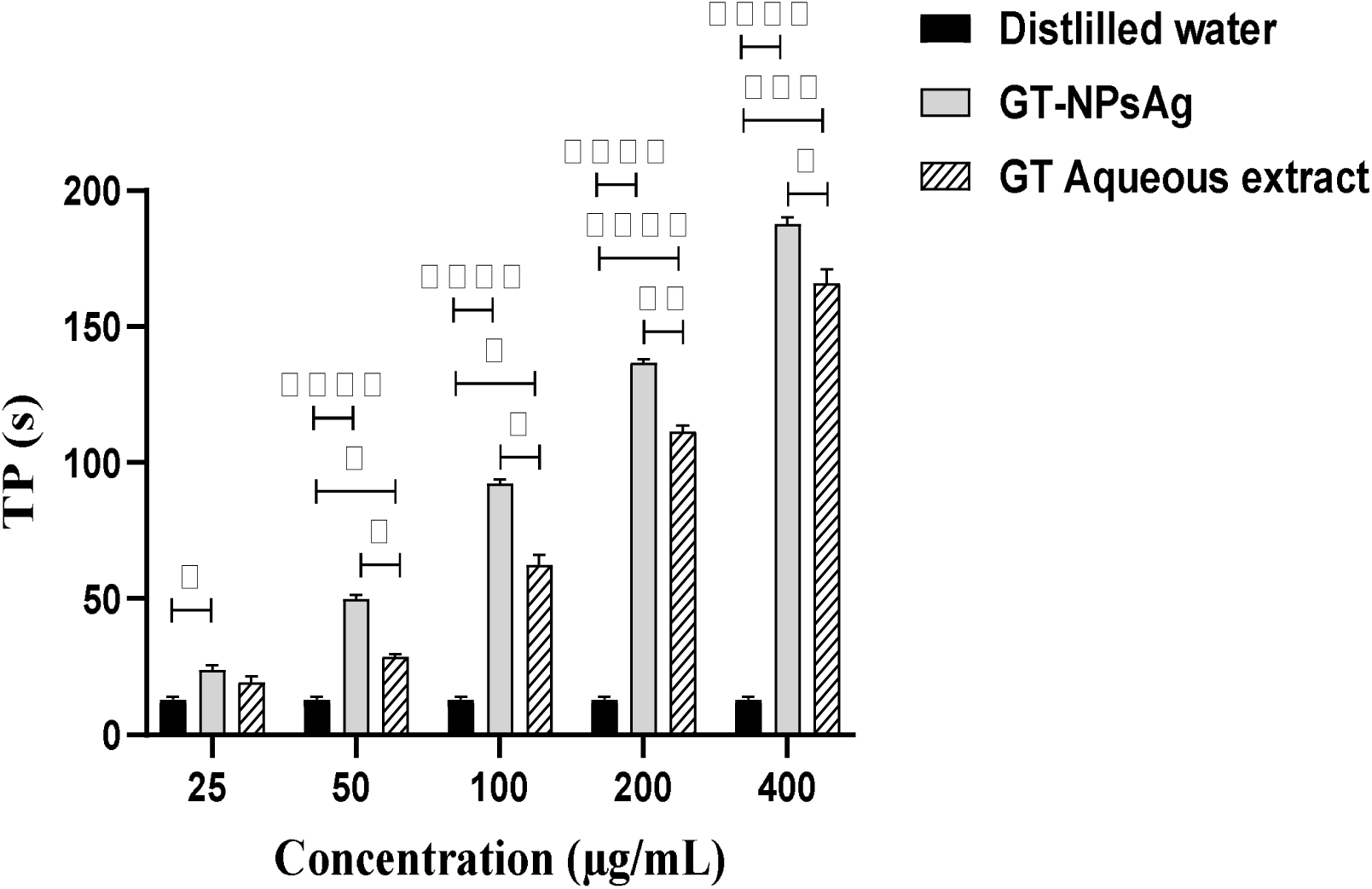
Anticoagulant activity on the exogenous coagulation pathway

The in vitro evaluation of the anticoagulant activity of the aqueous extract and *G. tessmannii*-mediated silver nanoparticles (GT-AgNPs) on the extrinsic coagulation pathway demonstrated that both preparations exhibited anticoagulant effects. The negative control displayed a clotting time of approximately 13 s. Comparison of the prothrombin time (PT) obtained at different concentrations (25, 50, 100, 200, and 400 μg/mL) showed that the highest concentration (400 μg/mL) produced the greatest anticoagulant effect. At this concentration, GT-AgNPs prolonged PT to 188 s, corresponding to an extension of 175 s compared with the negative control, whereas the aqueous extract produced a PT of 166 s, representing a prolongation of 153 s relative to the negative control. These results indicate that GT-AgNPs exert a stronger anticoagulant activity than the crude aqueous extract.

## IV. Discussion

Phytochemical analysis conducted in this study revealed that the aqueous bark extract of *Guibourtia tessmannii* is rich in phenolic compounds, particularly flavonoids, tannins, anthraquinones, and saponins. These findings are consistent with previous reports describing G. tessmannii as an abundant source of polyphenols, especially flavonoids. [32]. Such metabolites are widely recognized for their antioxidant, anti-inflammatory, and anticoagulant activities, which may explain part of the biological effects observed in the present study [33, 34]. Silver nanoparticles (AgNPs) were successfully synthesized through the green reduction of silver nitrate mediated by secondary metabolites present in the aqueous extract. These phytochemicals serve a dual function by reducing Ag⁺ ions to metallic silver (AgC) while simultaneously acting as capping and stabilizing agents, thereby preventing nanoparticle aggregation [35]. The formation of AgNPs was first evidenced by the progressive color change of the reaction mixture from light brown to dark brown after 48 hours of incubation, indicating nanoparticle formation. This observation was further confirmed by UV–Visible spectroscopy, which revealed a characteristic surface plasmon resonance (SPR) band at 450 nm, reflecting the nucleation and growth of silver nanoparticles [36]. Similar observations were reported for *Megaphrynium macrostachyum*, where incubation with silver ions produced a rapid color change accompanied by an SPR band between 400 and 550 nm [36]. The pH of the reaction medium has previously been shown to play a crucial role in AgNP biosynthesis, with alkaline conditions favoring the production of smaller, highly stable, and monodisperse nanoparticles [35, 36]. Likewise, the concentration of silver nitrate markedly influences nanoparticle formation. Song and Kim (2009) demonstrated that increasing AgNO₃ concentrations from 0.1 to 2 mM significantly improved nanoparticle stability, with 2 mM producing the most stable colloidal suspension [37]

Powder X-ray diffraction (PXRD) analysis revealed four characteristic diffraction peaks corresponding to the (111), (200), (220), and (311) crystallographic planes of face-centered cubic (fcc) metallic silver, confirming the crystalline nature of the synthesized nanoparticles. Application of the Scherrer equation yielded an average crystallite size of approximately 20 nm, which falls within the range commonly reported for plant-mediated silver nanoparticles characterized by PXRD [35].

Fourier-transform infrared (FTIR) spectroscopy further confirmed the involvement of hydroxyl-containing phytochemicals in nanoparticle synthesis. The observed shifts in characteristic absorption bands indicate that phenolic hydroxyl groups participated in both the reduction of Ag⁺ ions and the stabilization of the resulting nanoparticles. These findings agree with those of Guo et al. (2025), who demonstrated the important role of phenolic compounds in the green synthesis and stabilization of silver nanoparticles [38].

Energy-dispersive X-ray spectroscopy (EDX) detected silver, carbon, and oxygen, confirming the presence of organic molecules adsorbed on the nanoparticle surface. Scanning electron microscopy (SEM) further revealed aggregated nanoparticles exhibiting irregular fibrous-like morphologies. Comparable structural features have been reported for silver nanoparticles synthesized from Coriandrum sativum and Selaginella extracts [39].

Acute oral toxicity studies demonstrated no evidence of treatment-related toxicity following administration of GT-AgNPs at the OECD limit dose of 2000 mg/kg body weight. Clinical observations, body weight gain, organ-to-body weight ratios, and biochemical markers of hepatic and renal function remained largely unchanged compared with the control group.

Although serum AST activity showed a statistically significant increase, the measured values remained within normal physiological limits as reported by Klein (2021) [40]. No mortality was recorded at the 2000 mg/kg limit dose, indicating that the LD50 of GT-AgNPs exceeds 2000 mg/kg. These findings are consistent with those of Shanker et al. (2017), who reported no toxicity to the heart, liver, or pancreas following administration of silver nanoparticles derived from *Psoralea corylifolia* at the same dose [41]. Similarly, Songue Choudhary *et al.* (2022) observed no signs of liver or kidney damage and no mortality in rats treated with GT-AgNPs at 2000 mg/kg [42].

Evaluation of erythrocyte compatibility demonstrated that both the aqueous extract and GT-AgNPs exhibited very low, dose-dependent hemolytic activity, with hemolysis rates remaining below 10%. According to accepted hemocompatibility criteria, such values indicate that the tested materials are non-hemolytic. Similar observations were reported by Obiang et al. (2021) for aqueous extracts of G. tessmannii and by Dakshayani et al. (2019) for Selaginella-derived silver nanoparticles [20, 43].

The anti-inflammatory activity was first evaluated *in vitro* using the heat-induced egg albumin denaturation assay. Both the aqueous extract and GT-AgNPs inhibited protein denaturation in a concentration-dependent manner, with GT-AgNPs producing the highest inhibition (97.71%) at 400 μg/mL, exceeding both the aqueous extract and the reference drug diclofenac. These findings suggest that nanoformulation enhances the anti-inflammatory potential of the phytochemicals, probably by increasing their stability and improving interactions with protein targets.

The in vivo anti-inflammatory activity was assessed using the carrageenan-induced paw edema model, a well-established model of acute inflammation comprising three sequential phases involving histamine and serotonin release (first hour), kinins (second hour), and prostaglandins together with cyclooxygenase products (third phase) [44]. GT-AgNPs significantly reduced paw edema throughout the inflammatory response, with the greatest inhibition (94.92%) observed 5 hours after administration of 0.4 mg/kg body weight. These results are consistent with previous studies demonstrating enhanced anti-inflammatory activity of phytosynthesized silver nanoparticles at relatively low doses [35]. The superior activity of GT-AgNPs compared with the aqueous extract suggests that nanoparticle formation enhances the bioavailability and biological efficacy of the phytochemicals.

To investigate their anticoagulant properties, platelet-poor plasma (PPP) was used to evaluate both intrinsic and extrinsic coagulation pathways. GT-AgNPs prolonged both activated partial thromboplastin time (aPTT) and prothrombin time (PT) in a concentration-dependent manner, demonstrating anticoagulant activity through both coagulation pathways. At 400 μg/mL, GT-AgNPs produced the greatest prolongation of coagulation time, although their activity remained slightly lower than that of the reference anticoagulant enoxaparin.

Similar anticoagulant effects have previously been reported for silver nanoparticles synthesized from Selaginella species, which significantly prolonged plasma clotting time [43]. Collectively, these findings indicate that green-synthesized GT-AgNPs combine potent anti-inflammatory and anticoagulant activities, making them promising candidates for the management of thrombo-inflammatory disorders.

## V. Conclusion

This study investigated the anti-inflammatory and anticoagulant potential of silver nanoparticles biosynthesized from the stem bark extract of *Guibourtia tessmannii*. The work encompassed the green synthesis, physicochemical characterization, safety evaluation, and biological assessment of the resulting nanoparticles.

Silver nanoparticles were successfully synthesized using an aqueous extract of *G. tessmannii* through an environmentally friendly, cost-effective, and reproducible green synthesis approach. Characterization by UV–Visible spectroscopy, PXRD, FTIR, and SEM-EDX confirmed the formation of stable, phytochemical-capped silver nanoparticles with an average crystallite size of approximately 20 nm and an aggregated surface morphology.

Acute oral toxicity studies demonstrated a favorable safety profile, with no mortality or clinically relevant toxic effects observed following administration of the OECD limit dose of 2000 mg/kg body weight, indicating an LD₅₀ greater than 2000 mg/kg.

Biological investigations revealed that GT-AgNPs exhibited marked anti-inflammatory activity, significantly outperforming both the crude aqueous extract and the reference anti-inflammatory drug in the experimental models employed. Furthermore, the nanoparticles displayed dose-dependent anticoagulant activity on both the intrinsic and extrinsic coagulation pathways, while remaining non-hemolytic, demonstrating good blood compatibility.

The findings demonstrate that green-synthesized silver nanoparticles derived from Guibourtia tessmannii considerably enhance the biological activities of the parent plant extract, particularly with respect to anti-inflammatory and anticoagulant effects. These results support the potential application of GT-AgNPs as novel nanophytopharmaceutical candidates for the treatment of thrombo-inflammatory disorders. Future studies should investigate their molecular mechanisms of action, pharmacokinetic behavior, long-term safety, and therapeutic efficacy in disease-specific preclinical models to facilitate clinical translation.

## Ethics Statement

The animals were examined and adapted to the new environmental conditions for a week before the formal experiment. All experimental procedures were in strict compliance with the approved protocol by the Institutional Ethic Committee for Human Research of the University of Douala (Protocol approval number 3558CEI-UDo/03/2023/T).

## Data availability

the authors declare that the data supporting the findings of this study, including raw data files, are available from the corresponding author upon reasonable request.

